# The Effect of Proportion and Degree of Resistance of CD4^+^ Lymphocytes on in vivo HIV dynamics

**DOI:** 10.1101/466672

**Authors:** Andrew O. Cole, Rachel W. Mbogo, Livingstone S. Luboobi

## Abstract

For a disease like HIV(Human Immunodeficiency Virus) that infects 240 human beings each hour with the consequent increase in morbidity and mortality, a cure would be very welcome news. In order for this to happen, the true in vivo dynamics of transmission must be understood. Mathematical modelling and computational prowess have made it possible to convert large sets of data into easily analyzed and interpreted information with an amazing potential to change lives. In this paper, we explored the use of biomathematical modeling with computation in the dynamic changes resulting from HIV infection of CD4^+^(Cluster of differentiation 4) lymphocytes. This model is designed to simulate the in vivo dynamics of the proportion and degree of CD4^+^ lymphocyte resistance. Our results show that the degree of intrinsic CD4^+^resistance against the acquisition of HIV is more important than the proportion of resistant CD4^+^lymphocytes in vivo. Genetic determinants of resistance are postulated to be the backbone of development of intrinsic resistance. This work may serve as serve as a blueprint on how to use biomathematical modeling to tackle other infectious diseases. The findings of this work will be the basis of enthusiastic research to finally realize definitive control of HIV.

## 1. Introduction

### 1.1 Background

The burden of HIV on the human race has been hefty in terms of disease morbidity, mortality, and even disruption of family dynamics due loss of gainful work, increased orphan rates and financial strain on whole country economies from care of those affected. The loss of human labor, typically at the most productive ages of life, has also caused a synergistic effect culminating in the augmented economic burden to countries (“Global HIV targets,” 2015).

By the end of the year 2017 36.9 million people were living with HIV around the globe, a metric that obviates the urgent need to get this disease under control (“WHO ~ Data and statistics,"). Much has been done in the recent past around prevention of disease acquisition, progression limitation, and eradication. Although the current state of healthcare is devoid of a cure, treatment modalities that allow infected people to lead otherwise normal lives are fairly advanced (Chun, Moir, & Fauci, 2015). We have drugs that can restrict viral entry into *CD4^+^* cells, inhibit the functionality of reverse transcription, viral DNA (Deoxyribonucleic acid) integration into host DNA, even assembly and exit inhibition (De Clercq, 2009). Much yet remains to be done if HIV is to have a cure or a definitive preventative solution.

The mathematical model proposed and solved in this paper describes the in vivo dynamics of HIV transmission along with the most profitable areas of focus that could lead to a disease free equilibrium (DFE).

### 1.2 HIV Life Cycle

The life cycle of this disease begins when an HIV virion comes into physical contact with any human cell that expresses a cluster of differentiation type 4 (*CD4^+^)* on the surface of its cell membrane (Nicola et al., 2002). The cells that typically express these surface receptors are known as *CD4^+^* lymphocytes. This contact between an HIV virion and a *CD4^+^* lymphocyte needs another receptor (co-receptor) that aids viral entry into the host lymphocyte cytoplasm. These receptors are of 2 types, CCR5 (C-C chemokine receptor type 5) and CXCR4 (C-X-C chemokine receptor type 4), and together with the *CD4^+^* receptor form the perfect facilitation of viral entry. Besides lymphocytes, the other cells that express the *CD4^+^* receptors include tissue macrophages, dendritic cells, and gastrointestinal epithelial cells (Namazie et al., 1990).

Once the virion has gained entry into the host cytoplasm via a process called endocytosis, the virion uncoats from its envelope releasing its nucleic acid contents (RNA) (McMichael & Rowland-Jones, 2001). At this point the virus performs a reverse transcription reaction which produces provial DNA using the viral RNA as a template for the reaction and a reverse transcriptase enzyme to catalyze this reaction. The reverse transcriptase enzyme is contained in a gene (pol) which is of HIV origin (Charneau et al., 1994). With this step successfully completed the newly synthesized proviral DNA is inserted into the host DNA via integration by another enzyme called integrase (Bushman, Hansen, & Butler, 2001). This is a critical step because, after this process, the only way to remove or destroy viral DNA is by destroying the entire cell that contains this integrated DNA.

As host DNA replication is taking place, viral DNA will be replicated along with that of the host. The resultant mRNA and protein synthesis that takes place in the host cytoplasmic space produces new virions alongside the human host proteins. These undergo assembly into fully matured virions that can now exit the CD4^+^ expressing lymphocyte by budding or even as a result of bursting of the lymphocyte via a process known as lysis (Chen, Walker, Baltimore, Collins, & Kalams, 1998). The new virions are now ready to come into contact with, and infect other host cells that express *CD4^+^* on their cell membrane surface, thereby repeating the process.

### 1.3 RNA Sequencing

This is the processes by which the type and quantity of RNA is measured in vitro. RNA is very unstable (labile) and is normally only present in the system as the need for it arises. Once it has been used to attain certain cellular goals, it is quickly degraded (Levin et al., 2010). This means that the concentration as well as the type of RNA that is being expressed in the cell is in constant dynamism. The resultant challenge, unlike DNA (very stable), is that of accurately measuring how much RNA there is in the cell at any given time (t). The same problem applies to the type of RNA being expressed at any given time (t).

RNA sequencing is one answer in that it is a technology with the great promise, of accurately relaying qualitative as well as quantitative values of RNA in the cell.

The process begins with blood being collected in a container that keeps RNA viable and stable in order to “freeze” it in the exact proportions and varieties that are found in vivo (Bhagwat et al., 2003). This is then amplified in the RNA sequencing machine and bases are called at each position resulting in the RNA strand as it is in the cell (Levin et al., 2010).

This information is then cleaned and processed with the results ready for analysis. The end result is that, it is now possible to describe the true picture of the molecular dynamics happening intracellularly at any given time point.

The data analyzed for this manuscript was produced by RNA sequencing after *CD4^+^* cells were isolated, collected, and subjected to three different external conditions of lymphocytes stimulation artificially:

i. Unstimulated
ii. Stimulated for 4hrs
iii. Stimulated for 24hrs

The stimulation of *CD4^+^* lymphocytes is the artificial mimicry of in vivo realities in that (Wang et al., 2004), whole cells are stimulated in an in vitro setting and then later subjected to RNA sequencing to see if there are any changes in quality, quantity, peak concentration, gradient of concentration increase or decrease, and even related co-expression of genes. With this, we are able to see which subjects possess resistant cells and which subjects have the wild-type variety of cells. We also see what the response (reaction) of the cells is, when infection happens or when treatment is commenced. Just by looking at stimulated cells, otherwise infinitesimal changes become augmented enough for the changes to be appreciable.

### 1.4 Explanation of the potential effect of proportion and degree of, susceptible CD4^+^ lymphpocyte resistance, on resistance against HIV Acquisition

The proportion and degree of resistance represent a bivariate method of explaining how different individuals have varied susceptibility and resistance profiles (Cole, 2017). We postulate that this is informed, in large part, by the genetic basis of expression in each individual. Past studies have supported the genetic basis of resistance to HIV infection (“The Geographic Spread of the CCR5 Δ32 HIV-Resistance Allele,”) and we characterize the magnitude of such resistance profiles. To describe these influences on in vivo transmission dynamics of HIV, we have developed a graphical representation of a few possible scenarios.

In Figure 3, the Y-axis represents the proportion of resistant *CD4^+^* lymphocytes as compared to the proportion of *CD4^+^* cells that are wild-type. This axis only deals with the numbers of resistant cells as compared to those that have a wild-type genotype. There is no information measuring the magnitude of any given *CD4^+^* lymphocyte, meaning, every lymphocyte whose resistance profile lies above a predefined threshold will be considered resistant. Since we assume that every susceptible *CD4^+^* lymphocyte in the host is uniquely different in terms of resistance magnitude, we purposed to also measure the degree of resistance of any given susceptible but resistant *CD4^+^* lymphocyte. That magnitude of resistance is captured on the X-axis and will be modeled along with the dynamics of transmission.

**Figure 1:**
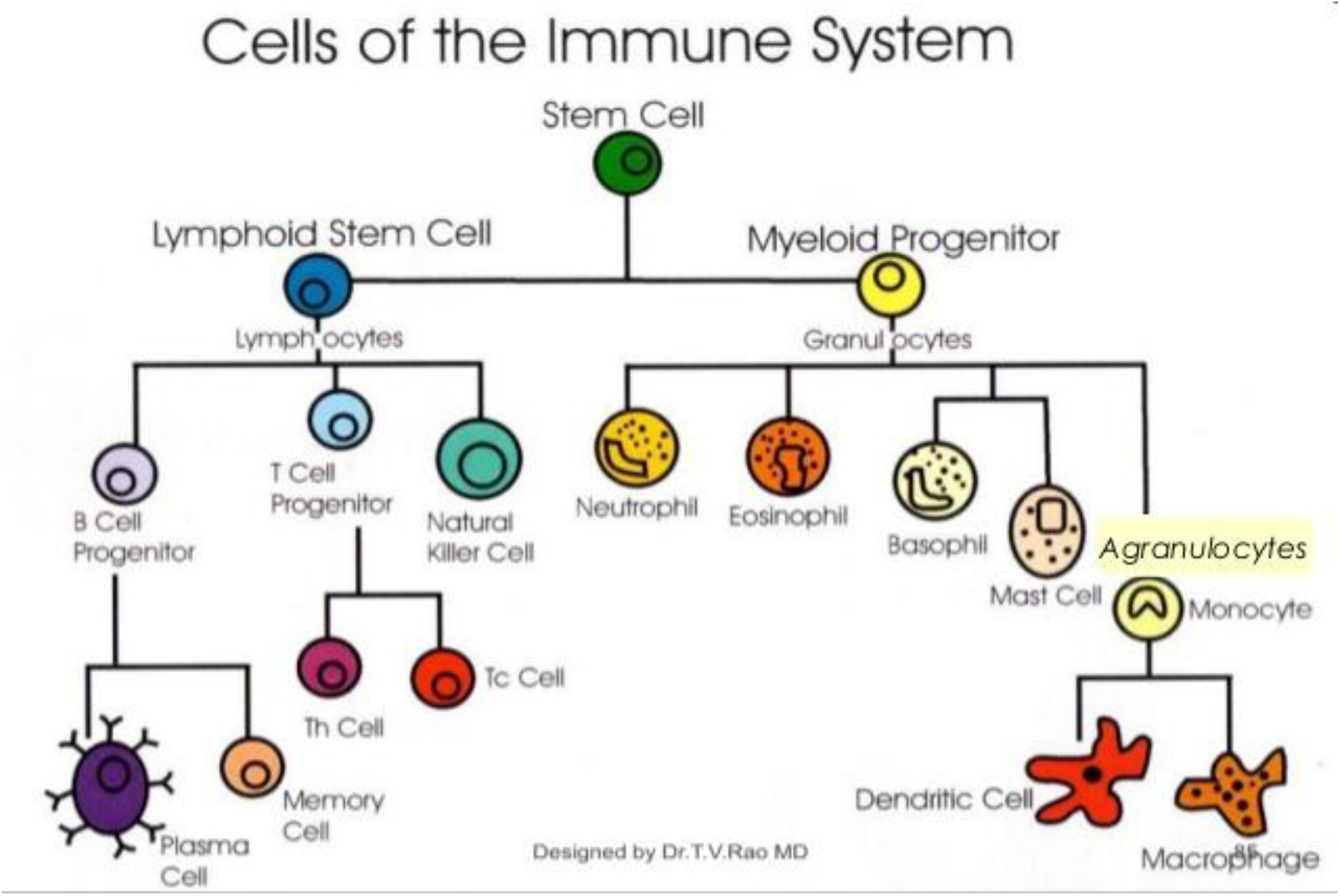
Cells of the immune system. https://image.slidesharecdn.com/macrophagesdendriticcellsinviralinfection-140109133413phpapp02/95/macrophages-dendritic-cells-in-viral-infection-9-638.jpg?cb=1389274654

**Figure 2:**
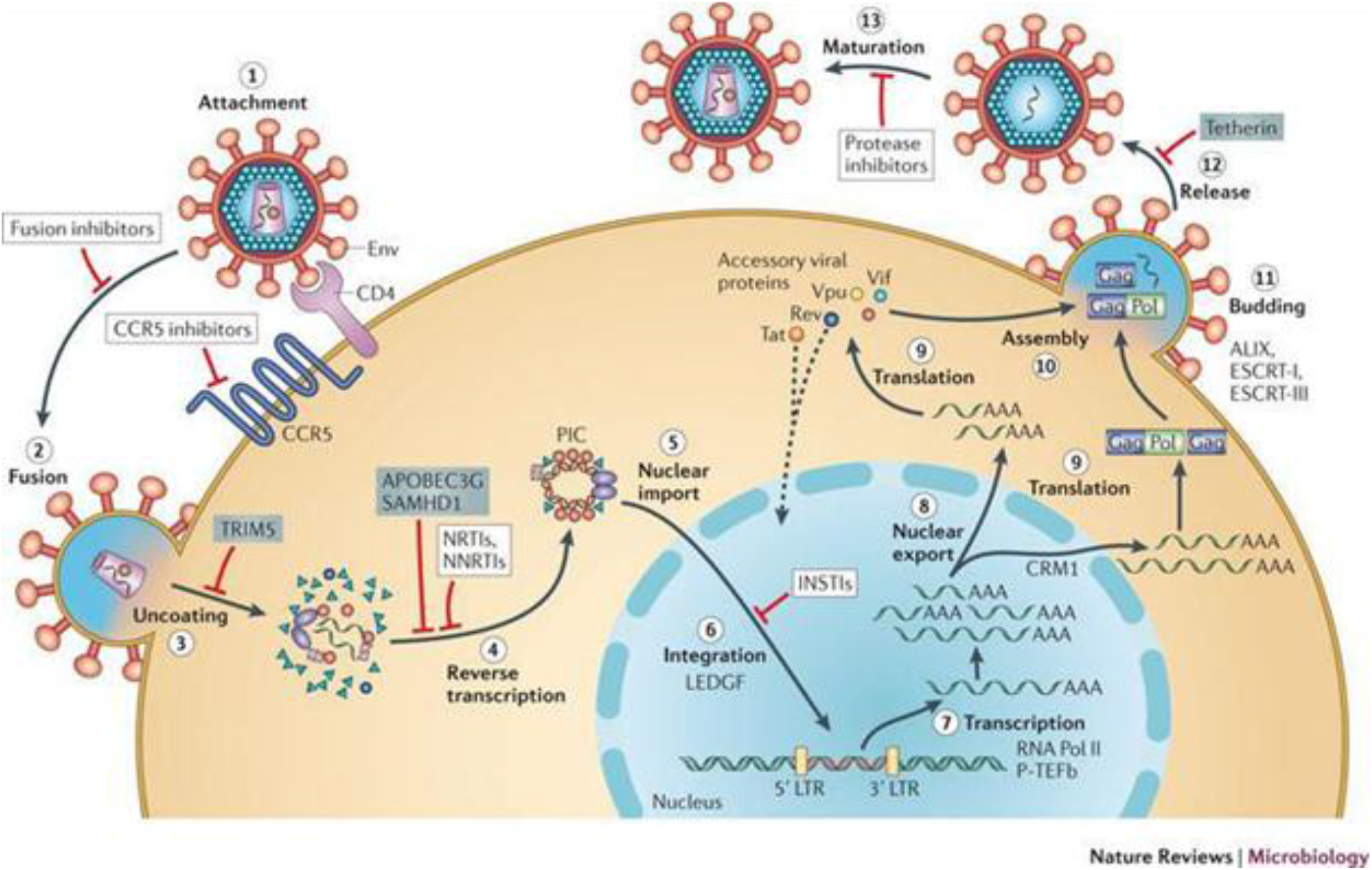
HIV life cycle. https://www.bing.com/images/search?view=detailV2&ccid=MvC%2bxT74&id=C951C3A8B83F715BB01D6DE603DD28DCD21AC5ED&thid=OIP.MvCxT74Sx9K5Kgl6ShDgAHaD_&mediaurl=https%3a%2f%2f294305267s7hqfks2cfh08ipwpengine.netdnassl.com%2fwpcontent%2fuploads%2f2016%2f02%2fhiv_gilead_antiretroviral_grant_aids_funding_research.jpg&exph=509&expw=946&q=hiv+life+cycle&simid=608028644288825368&selectedIndex=21&ajaxhist=0

**Figure 3:**
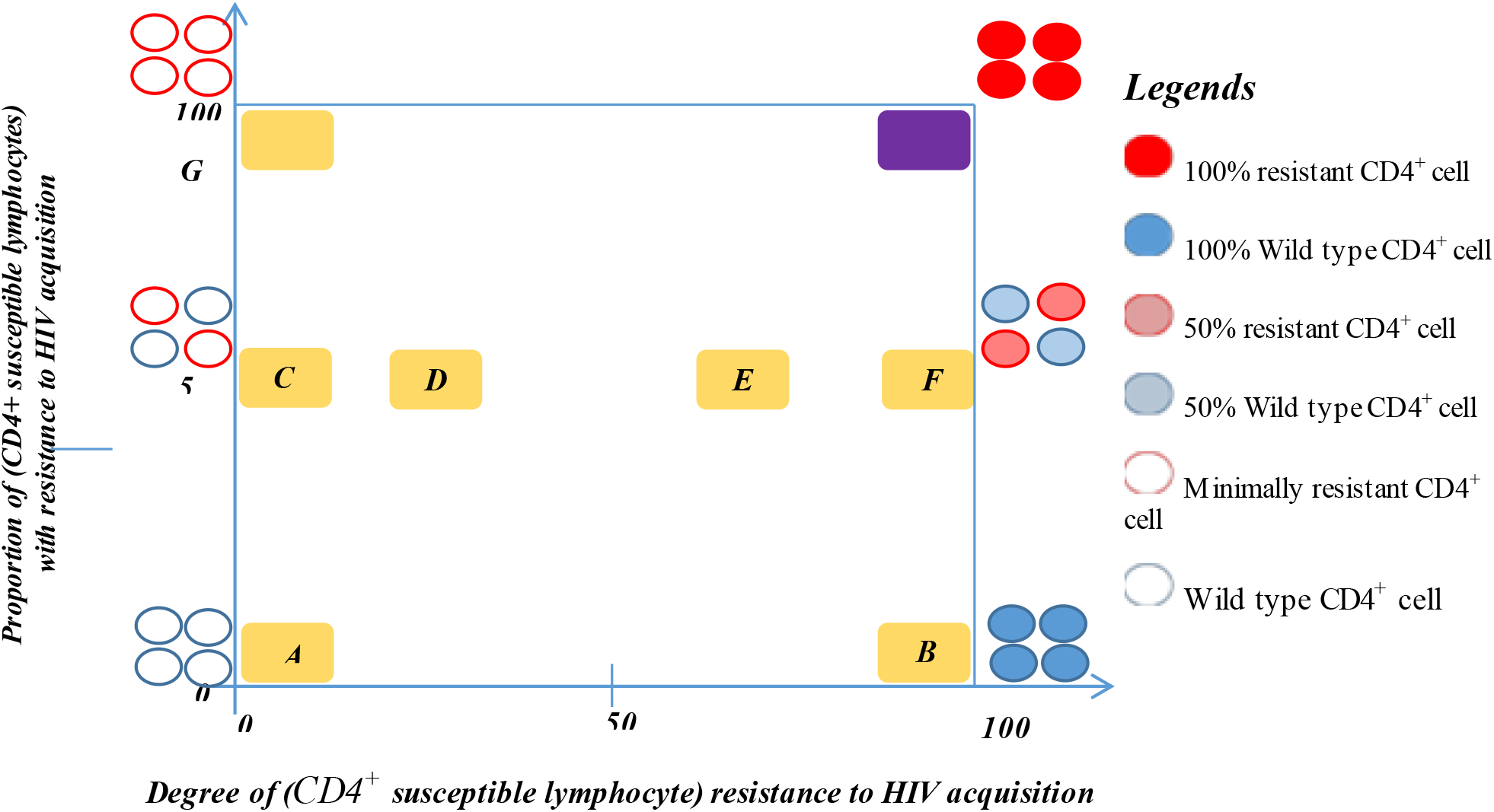
Proportion versus degree of susceptible *CD4^+^* lymphocytes resistance to HIV acquisition.

Several coordinates on Figure 3, representing the relationship between resistance (proportion and degree) versus transmission will be identified and simulated to show the effect in terms of in vivo dynamics.

## 2. Mathematical Model

### 2.1 Background

A robust predictive model for the acquisition of HIV in exposed groups is presented in this section. The section is organized with the assumptions for the in vivo dynamics of HIV and its interactions with cells of the immune system presented first. The conceptual diagram is presented next with a description of the state variables and parameters representing the interactions modeled and used in the analysis to reveal the most appropriate target of intervention (preventative or treatment) that would culminate in the biggest impact on HIV control. The sequential organization continues with a display of the system of ordinary differential equations describing all transmission dynamics as presented in the conceptual diagram. The analytical solution of the system of ODEs (Ordinary Differential Equations) is then presented next and this is followed by a section where the basic reproductive number and the effective reproductive number are derived. The logical progression of analyzing the system for local stability, and finding an endemic equilibrium (EE) as well as disease free equilibrium (DFE) concludes this section.

In order to demonstrate the applicability of this mathematical model, simulations based on changing the initial conditions, of state variables and specified values of parameters, are applied to the system of ODEs and displayed in Section 3.

### 2.2 Dynamics of HIV transmission

The dynamics of HIV transmission are governed by the following assumptions:

i. Lymphocytes are recruited at different rates from the germinal centers;
ii. The cause of a difference in the recruitment rates is due to varied lymphocyte simulation;
iii. Susceptible lymphocytes die naturally at fixed rate (1/lifespan);
iv. Lymphocytes can only be infected after interacting with the virus;
v. Infected lymphocytes die at different rates from that of susceptible lymphocytes;
vi. The rate of lymphocyte infection depends on the expression genetics of the susceptible lymphocyte;
vii. Infected lymphocytes do not recover;
viii. Age of the host affects the dynamics of in vivo lymphocyte infection; and host co-morbidity affects the rates of infection and the rates of death of lymphocytes.

### State variables and Parameters

The state variables and parameters in the model for the in-host HIV dynamics are summarized in Tables 1 and 2, respectively.

**Table 1:** The State Variables.

| **State Variable** | **Description** |
| --- | --- |
| $L_W$ | Number of Wild type susceptible lymphocytes per $\mu\text{l}$ (Microliter) |
| $L_R$ | Resistant susceptible lymphocytes per $\mu\text{l}$ |
| $V$ | Virus (free HIV virions) per $\mu\text{l}$ |
| $L_i$ | Infected lymphocytes per $\mu\text{l}$ |

**Table 2:** The Parameters.

| Parameter | Description |
| --- | --- |
| $\lambda$ | Recruitment rate of new $CD4^+$ lymphocytes |
| $p$ | Proportion of wild-type susceptible host lymphocytes ($CD4^+$ T cells) |
| $1 - p$ | Proportion of resistant susceptible host lymphocytes ($CD4^+$ T cells) |
| $\mu_w$ | Death rate of wild-type host lymphocytes ($CD4^+$ T cells) |
| $\mu_R$ | Death rate of resistant host lymphocytes ($CD4^+$ T cells) |
| $K$ | Number of virions released per dying/bursting infected $CD4^+$ lymphocyte |
| $\psi$ | Death rate of the HIV virion per unit time |
| $\mu$ | Natural death rate of an infected host lymphocyte ($CD4^+$ T cell) before releasing new virions |
| $\sigma$ | Death rate of an infected host lymphocyte ($CD4^+$ T cell) upon which new virions are released into the blood system |
| $\alpha$ | Rate of infection of the wild-type host lymphocytes by the virus |
| $\beta$ | Rate of infection of the resistant host lymphocytes by the virus |

These dynamics of HIV infection governed by the state variables and parameters in 2.2 are depicted in Figure 4

**Figure 4:**
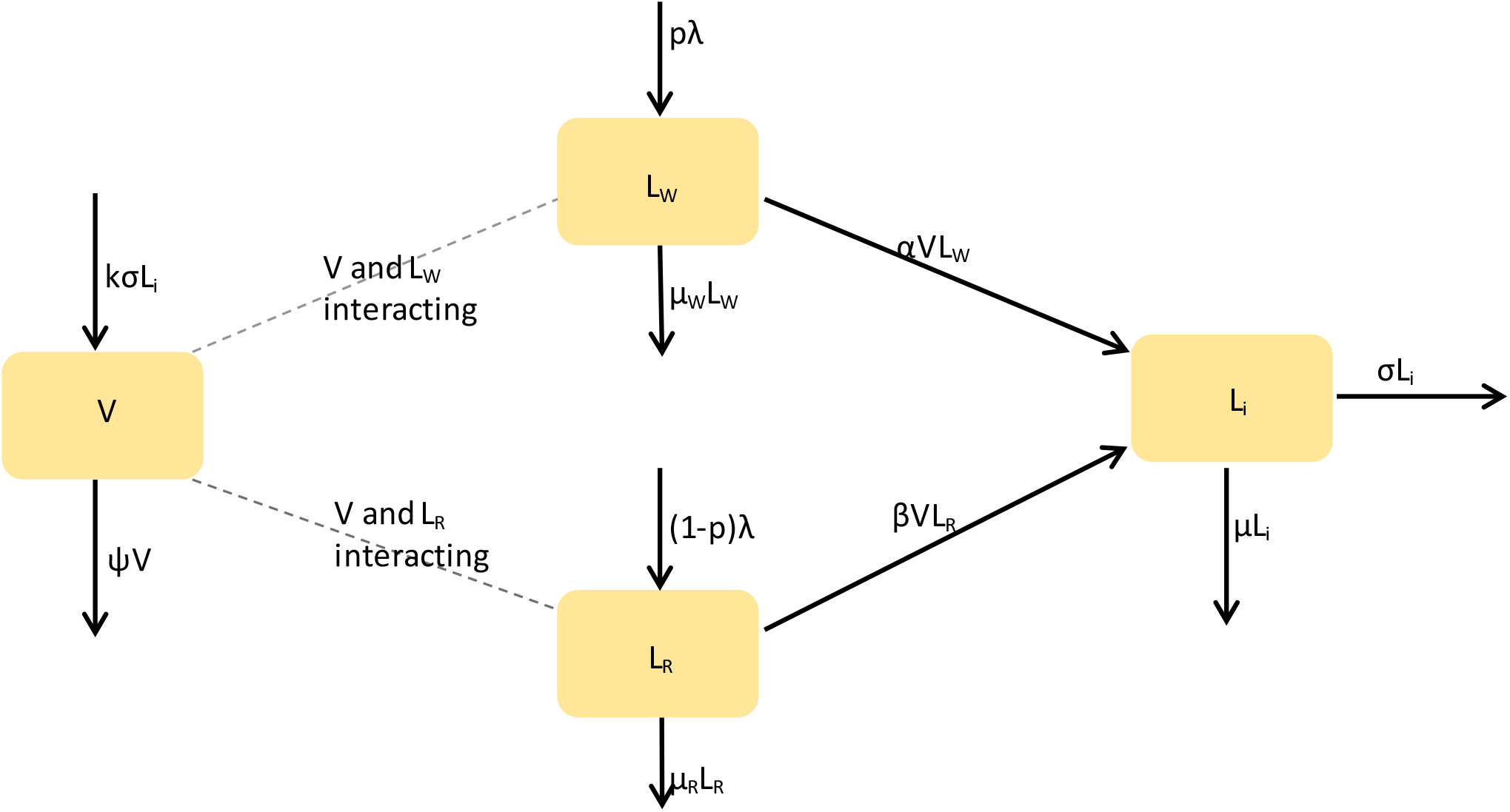
Compartmental diagram displaying In-host interaction between HIV virions and the Lymphocytes.

#### 2.2.1 System of Ordinary Differential Equations for the model

From the description and assumptions made regarding the dynamics of in vivo HIV transmission dynamics where the *CD4^+^* susceptible lymphocytes are either wild-type or resistant, as depicted in the compartmental diagram, we derive the model as in (M1):

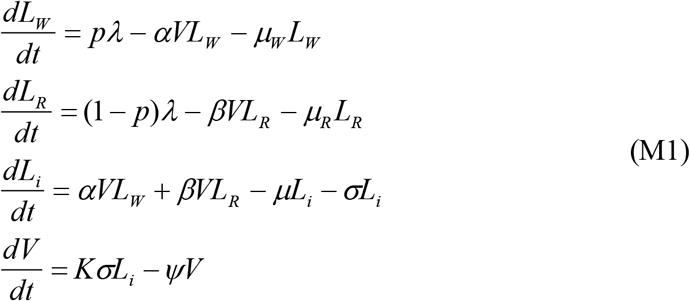

#### 2.2.2 Analysis of the Model

First we investigate the possibility of existence of equilibrium solutions of model (M1). At equilibrium the derivatives in (M1) are zero. That is:

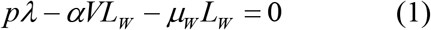

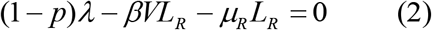

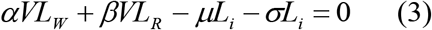

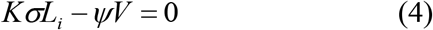

Suppose 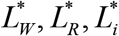 and *V*\* are the equilibrium values of the respective variables as derived from equations (1) – (4). Then we have:

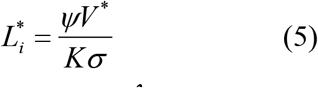

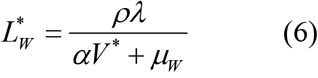

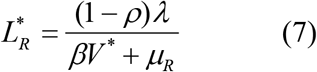

Thus for *V*\*= 0, we have 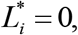 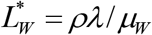 and 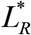 These are the disease free levels of the lymphocytes.

For we have from (1) and (2) we have:

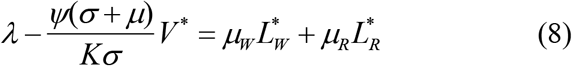

In terms of *V*\*, equation (8) becomes:

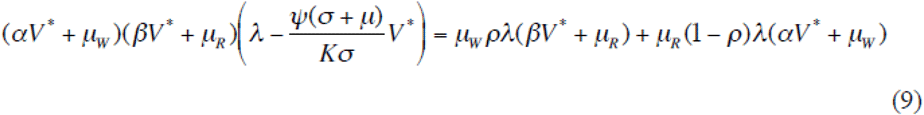

Equation (9) is cubic equation in terms of *V*\*. On investigation it is found that only non-positive values of *V*\* are feasible.

### Deduction

An equilibrium solution to model (M1) exists only if the HIV is at level zero. So once one gets infected the HIV persists at different concentrations, which may be influenced by therapy that could control the values of the parameters of the model. Thus it is important to investigate further the effects of the values of the parameters. In this manuscript we demonstrate this in Section 3 through simulations of model (M1).

## 3 Simulation of the Mathematical Model

In the following simulations, we use the initial conditions for state variables and parameter values as indicated in Table 3.

**Table 3 (a):** Possible initial values of the State Variables.

| State Variables | Possible initial Value | Citation |
| --- | --- | --- |
| $V$ | 1200/μL | Estimate |
| $L_W$ | 750/μL | Nowak et al. (1996) |
| $L_R$ | 750/μL | Estimate |
| $L_i$ | 100/μL | Estimate |

**Table 3 (b):** Possible parameter values.

| Parameters | Value | Reference: |
| --- | --- | --- |
| $k\sigma L_i$ | 5/pL(Picoliter) | Estimate |
| $\Psi$ | 2.4/pL | Wordaz and Nowak (1999) |
| $p$ | 10/μL.day | Nowak et al. (1996)<br>Rong, Gilchrist, Feng, & Perelson, (2007) |
| **1-p** | **10/μL.day** | Estimate |
| **$\mu_w$** | **0.02/pL.day** | Srivastava and Chandara (2008) |
| **$\mu_R$** | **0.02/pL.day** | Estimate |
| **$\alpha$** | **0.2** | Nowak et al. (1996b) |
| **$\beta$** | **0.24** | Estimate |
| **$\mu_{Li}$** | **0.24/pl.day** | Odhiambo et al (2015) |
| **$\sigma_{Li}$** | **0.24/μL** | Wordaz and Nowak (1999) |

The following publications and manuscripts informed the choice of the various values: Wodarz & Nowak (2000), (Nowak et al., 1996), Rong et al. (2007), Srivastava & Chandra (2010), Odhiambo, Mbogo, & Luboobi (2015)

There are four distinct types of simulations presented in this paper.

i. Simulations where all the parameters are kept constant while the initial ratio of wild-type to resistant lymphocytes is varied;
ii. Simulations where the initial ratio of wild-type to resistant lymphocytes is kept constant while the parameter values representing the force of infection is varied in sequence;
iii. Simulation with a combination of initial values (State Variables) and parameters are varied simultaneously;
iv. Simulation of iii) above with a lengthening of the time scale to 10 months, as illustrated in Section 3.1.

### 3.1 Changing the proportion of Wild-type versus resistant susceptible lymphocytes while keeping all parameters constant

#### 3.1.1 Scenario 1: Resistant = 0, Wild-type = 1.0, Virus = 1200, and Infected = 100

Graph (Fig. 5) shows the dynamics of four state variables (Virus, Wild-type susceptible lymphocytes, Resistant susceptible lymphocytes, and infected lymphocytes) when the proportion of genotypically resistant lymphocytes is zero (R = 0). This assumes a highly unnatural proposition where all lymphocytes are deemed to be genotypically wild-type and therefore easily infected by virus upon interaction. It is a simulation of a fairly homogenously sensitive susceptibility profile for the lymphocytes. The resultant graph in Figure 5

**Figure 5:**
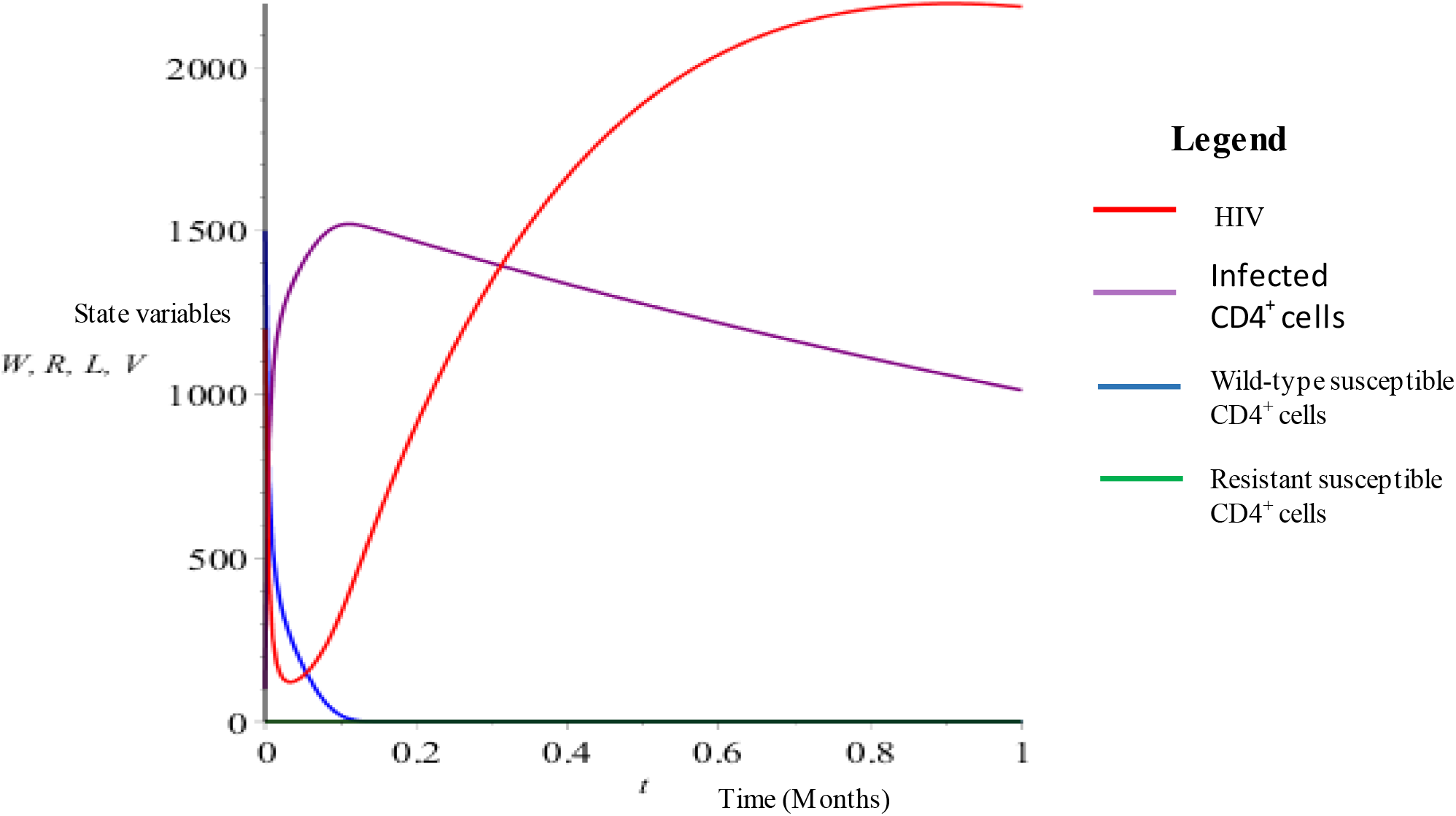
State variables vs time (no resistant lymphocytes)

shows only wild-type lymphocytes (blue curve) and an absolute lack of the resistant variety of susceptible lymphocytes (green curve-absent). Notice that each graph will have a total susceptible lymphocyte count of 1500 when the number of susceptible but wild-type lymphocytes are added to the number of susceptible but resistant lymphocytes. That is ***Total number of susceptible lymphocytes (1500) = number of Wild-type susceptible lymphocytes + number of Resistant susceptible lymphocytes.***

Since in this first scenario there are no resistant susceptible lymphocytes, all 1500 susceptible lymphocytes at initial conditions are wild-type. As time progresses the number of remaining susceptible lymphocytes drops in a relatively rapid fashion (blue curve) to almost complete depletion within 0.1 on the time scale while the number of infected lymphocytes (purple curve) shoots in an almost vertical way to reach its peak of near 1500 lymphocytes which are now infected. As time progresses past the 0.1 mark on the x-axis towards the 0.2 mark and beyond, the system is unable to make new susceptible lymphocytes to sustain the dynamics since all the wild-type susceptible lymphocytes have been depleted by seroconversion and now the infected lymphocytes begin dying off due to the shortened lifespan of seroconverted lymphocytes that has been shown to be 30 days as opposed to an average of 87 days. The population of virus in this in vivo system dynamics (red curve) initially drops extremely rapidly from an initial value of 1200 virions to approximately 120 virions then the curve reaches its trough where an inflexion point is realized as the viruses find and infect sufficient numbers of wild-type susceptible lymphocytes to eventually increase their numbers as the infected lymphocytes burst and release new virions in large numbers due the shortened lifespan of the now infected lymphocytes. The dynamics of the virus reach a maximum and again attain an inflexion point around the 0.8 mark on the time scale and will begin to drop as infected lymphocyte number is also depleted.

#### 3.1.2 Scenario 2: Resistant = 0.4, Wild-type = 0.6, Virus = 720, and Infected = 100

In vivo transmission dynamics when the initial conditions of the state variables are heterogenous but with a composition of 60% wild-type susceptible lymphocytes and 40% resistant susceptible lymphocytes produces the graphical output below (Fig. 6) when the initial condition of effective virus is also dropped by a similar proportion of 40% from the starting value of 1200. In this case, there will be 720 virions that have meaningful and potentially infection interactions with wild-type susceptible lymphocytes. The change in slope from more to less steep has happened sooner on the time scale because more viruses are interacting with resistant susceptible lymphocytes as opposed to the more efficient interaction between viruses and the wild-type variety of susceptible lymphocytes when considered from the viral point of view. The increase in “meaningless” interactions with resistant susceptible lymphocytes is inversely proportional to the more efficient interaction with wild-type susceptible lymphocytes. This, in essence, means that more viruses stay in the highly toxic environment of the blood stream and will be eliminated long before they get the chance to interact with the wild-type susceptible lymphocytes. This decrease in the number of available viable viruses allows the susceptible lymphocytes to persist in blood until past the 0.2 mark on the time axis, which is beneficial to the host that has this ratio of heterogeneity. It, however, is not the right heterogenous composition that will allow longevity of the host. It should be noted that unlike the first graph (Figure 5) where all susceptible lymphocytes carried a wild-type genotype, this heterogenous composition of 60:40 (wild-type: resistant) susceptible lymphocytes result in a drastic decline in virus load shortly after the simulation begins to a level of about 50 virions from the initial 720 virions. It is presumed that the virus has been able to establish and multiply due to its efficiency of infection upon eventual interaction with a wild-type susceptible lymphocyte.

**Figure 6:**
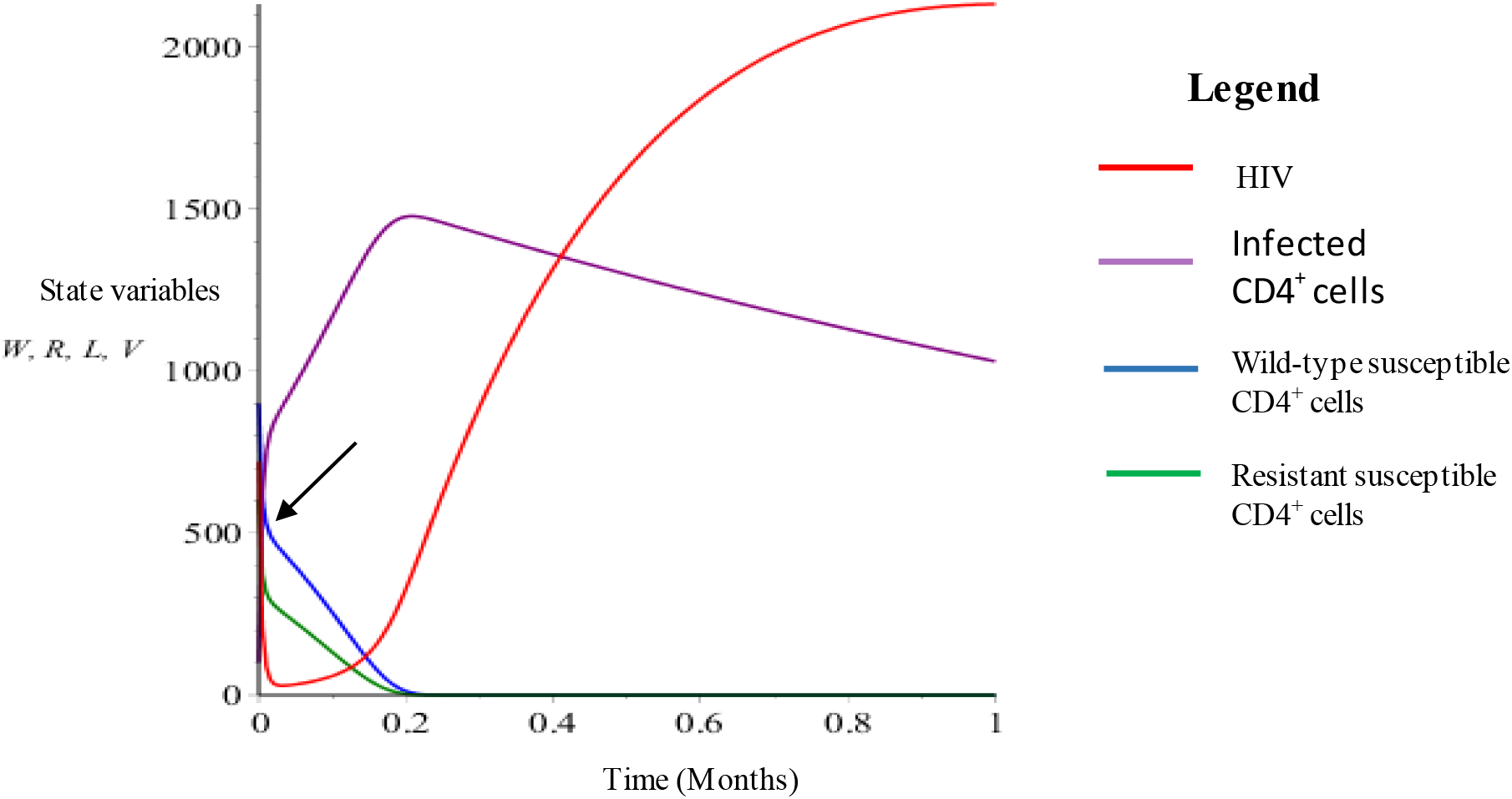
State variables vs time (40% resistant lymphocytes)

#### 3.1.3 Scenario 3: Resistant = 1.0, Wild-type = 0, Virus = 0, and Infected = 5

This Graph (Fig. 7) where the state variables are being sequentially altered at initial conditions shows the other extreme where another homogenous in vivo environment is the reality. Here we have 100% of the susceptible lymphocytes being resistant to HIV infection even when it interacts with an HIV particle. Although this scenario is not the most realistic, it offers the best protection to the host. As can be seen in Fig. 7, the resistant susceptible lymphocytes persist in the dynamic model for prolonged durations while the proportion of them that do seroconvert are few and they never really get to establish and propagate HIV. Free virus (red) will always interact with resistant susceptible lymphocytes resulting in massive elimination of virus. This means that the amount of effective infective virus is almost negligible. This is why the value given for virus at initial conditions is zero. The number of infected lymphocytes at initial conditions has also been halved to 5 because the in vivo ecosystem is so toxic for viruses that the eventual production of infected lymphocyte will be very low.

**Figure 7:**
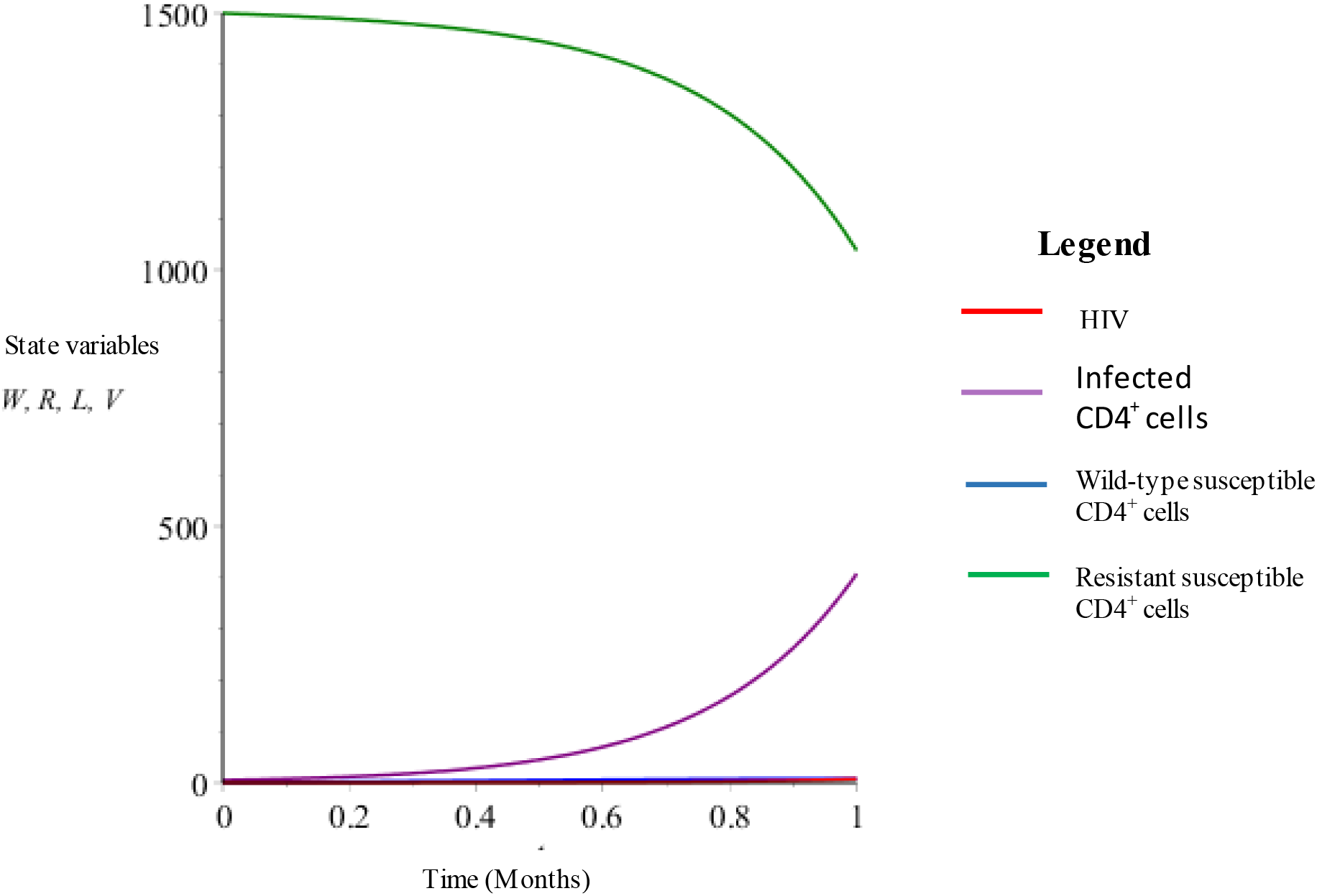
State variables vs time (100% resistant lymphocytes)

All the 3 Graphs (Fig. 5-Fig.7) in this Section (3.1) show that eventually even the resistant susceptible lymphocytes do get infected by virus. This could be considered a limitation of this model. This limitation has been taken into account in the next section where partial resistance is included in the model.

### 3.2 Changing the parameters of resistant lymphocytes while keeping all the initial conditions constant

This section contains for graphical output related to the dynamic transmission effect of changing only the transmission parameter of the resistant but susceptible lymphocytes and leaving the rate of transmission for the wild-type lymphocytes intact at initial conditions. Figure 8 displays what the dynamic simulation will look like as long as the proportion of lymphocytes initially is 1:1 between wild-type and resistant susceptible lymphocytes.

**Figure 8:**
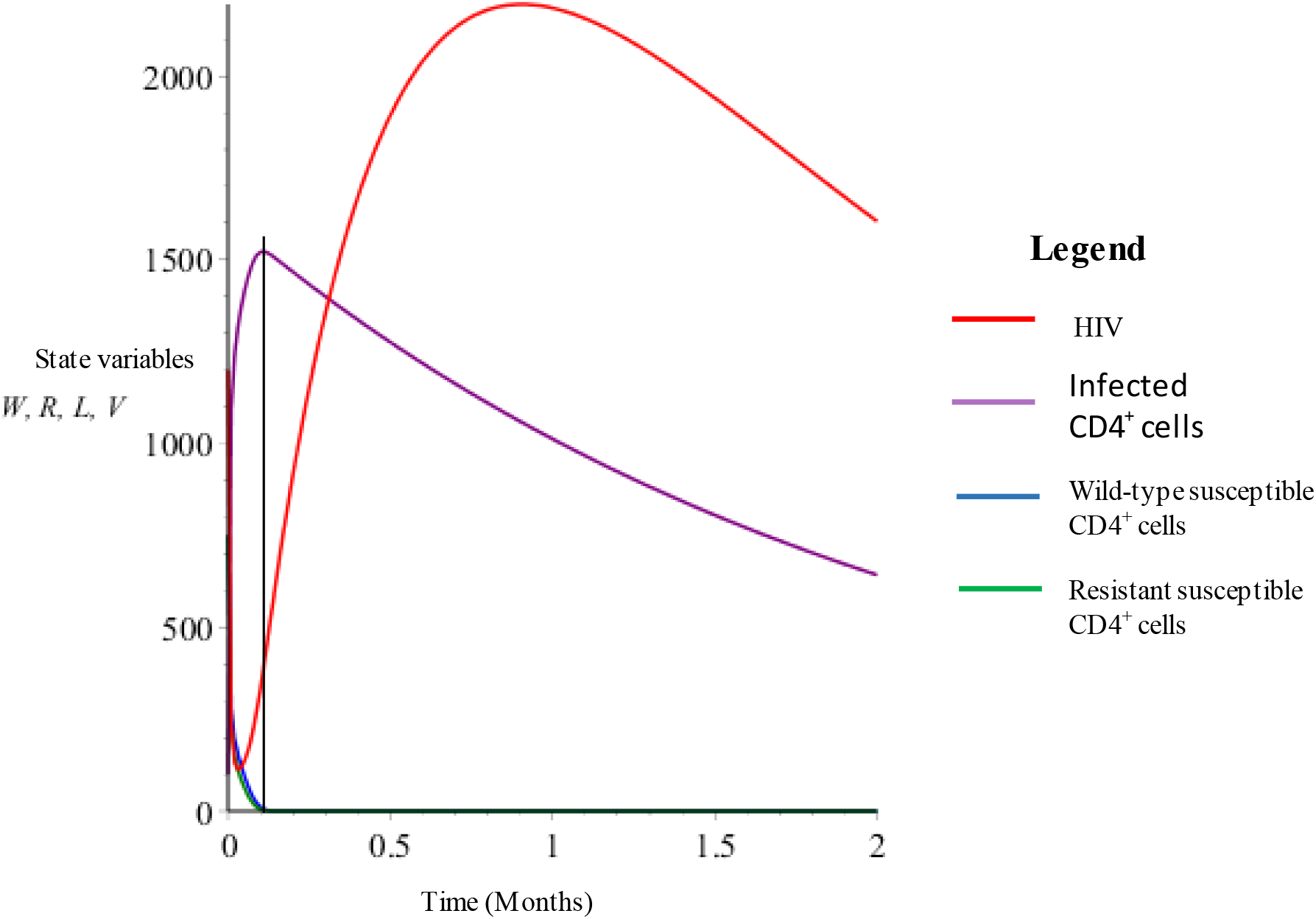
State variables vs time (β = 0.24 resistant lymphocytes)

#### 3.2.1 Scenario 1: β = 0.24, V = 1200, W = 750, R = 750, L = 100

As can be seen in Fig. 8, the wild-type (blue curve) and resistant (green curve) lymphocytes are almost superimposed over each other since the transmission rate (α and β) are similar (0.2 and 0.24) respectively. The dynamics show a rapid decline in both kinds of susceptible lymphocyte counts while a summation increase in infected lymphocytes is realized. This takes a very short time, as can be seen in the x-axis. The peak of the infected lymphocytes is reached in less than 2 months (purple curve) while the concentration of virus rapidly increases and reaches peak concentrations fairly early. Finally, as time progresses, the infected lymphocytes decline in number due to natural death as well as death from infection. The viral load (red curve) also starts to decline in tandem with the declining infected lymphocytes because there is a direct proportionality between infected lymphocyte counts and viral load. Notice that as long as the transmission rate of the resistant susceptible lymphocyte is similar in value to that of wild-type susceptible lymphocytes, the infection rate of lymphocytes is driven very rapidly reaching peak lymphocyte infection levels fairly early in the simulation progression. The entire system takes a short time to traverse each check point in HIV dynamics.

#### 3.2.2 Scenario 2: β = 0.01, V = 1200, W = 750, R = 750, L = 100

When all the state variables and parameters remain unchanged with the only change being in β (the transmission rate of resistant susceptible lymphcytes to β = 0.01), the transmission dynamics change. In this simulation (Fig. 9), the rate of transmission has been reduced in the resistant susceptible lymphocyte group to a value, 20 times less efficient than the wild-type susceptible lymhocytes. These wild-type susceptible lymphocytes are rapidly depleted (blue curve) because they are relatively easily transformed into infected lymphocytes after interaction with virus while the resistant susceptible lymphocytes take relatively longer to be depleted due to their markedly reduce rate if infection following interaction with viruses until they (blue curve) reach a value of near zero a full 6 months ahead of the more resistant lymphocytes. The infected lymphocytes show a curve with two obvious tangential slopes on the left and right of the black arrow. This is best explained by the fact that the virus is able to find (and infect) wild-type lymphocytes very efficiently yielding a curve with a steep slope. When most of the wild-type susceptible lymphocytes are depleted and most susceptible lymphocytes remaining are of the resistant variety, the tangential slope of the infected lymphocyte curve must be less steep owing to the less efficient transmission rate of HIV upon interaction with virus as compared to the previous interaction with wild-type lymphocytes. This delays the time (on the x-axis) before the peak level of infected lymphocytes is reached. This is advantagious to the lymphocytes involved because they get to live longer and therefore the host will by extension get to live longer too. The idea is therefore to make the resistant susceptible so resistant that the transmission rate to infected status is so slow that the virus either dies before infecting resistant but susceptible lymphocytes or the number of successfully infected, resistant, lymphocytes is so low and so slow that peak levels are never attained in a lymphocytes’ natural lifespan.

**Figure 9:**
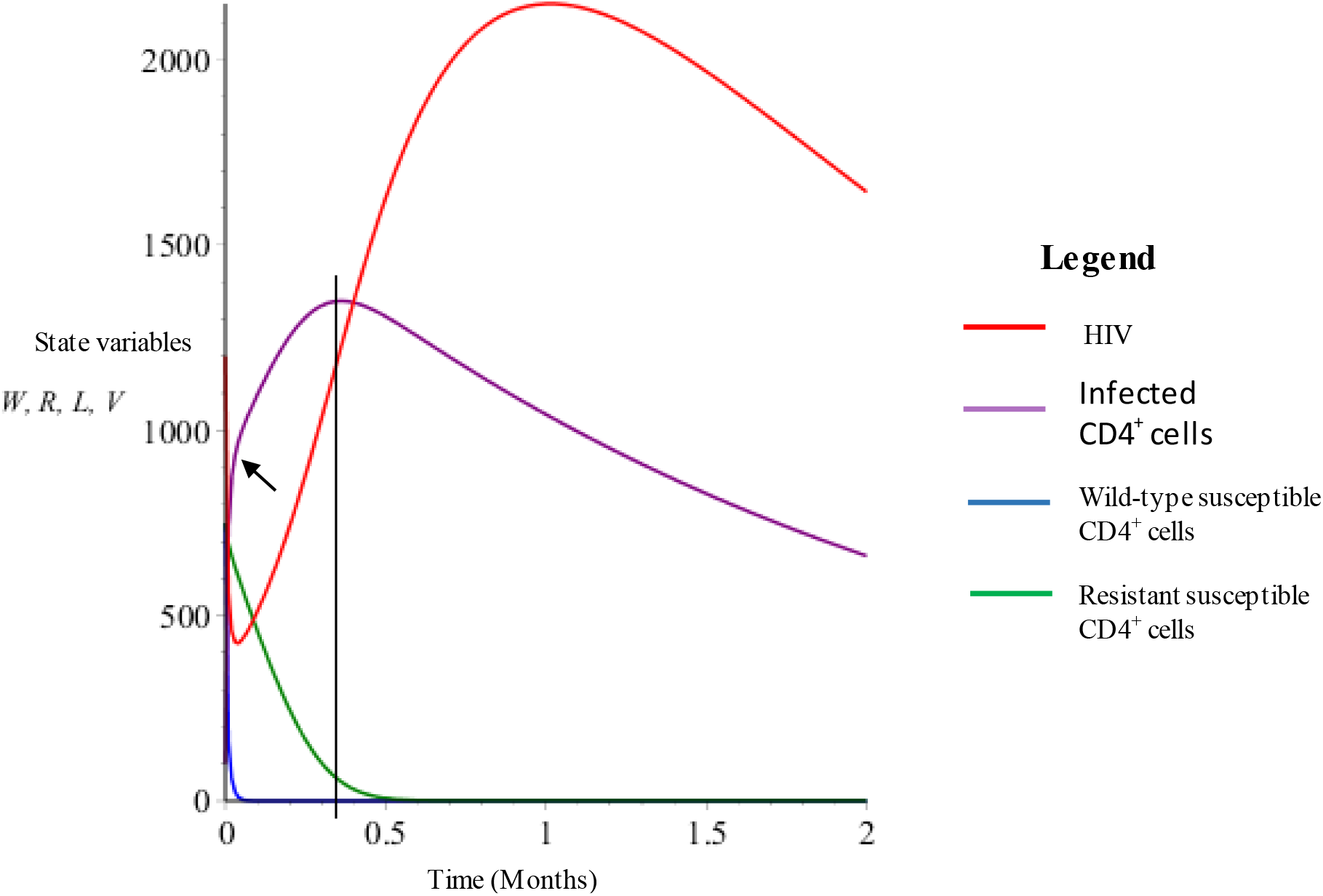
State variables vs time (β = 0.01 resistant lymphocytes)

#### 3.2.3 Scenario 3: β = 0.00001, V = 1200, W = 750, R = 750, L = 100

If the rate of infection in the resistant group is significantly less than that of the wild-type lymphocytes (20,000 times less efficient), the resultant Graph (Fig. 10) is depicted above. The wild-type lymphocyte count is depleted very rapidly, as has been seen previously, while the rate of decline of resistant susceptible lymphocytes (blue curve) now become negligible for more than two months. The transmission dynamics of the infected lymphocytes and virus (purple and red curves respectively) remain similar to prior graphs in trajectory although the rates are different. This is mainly because the real drive of transmission dynamics in this scenario is only the wild-type susceptible lymphocytes.

**Figure 10:**
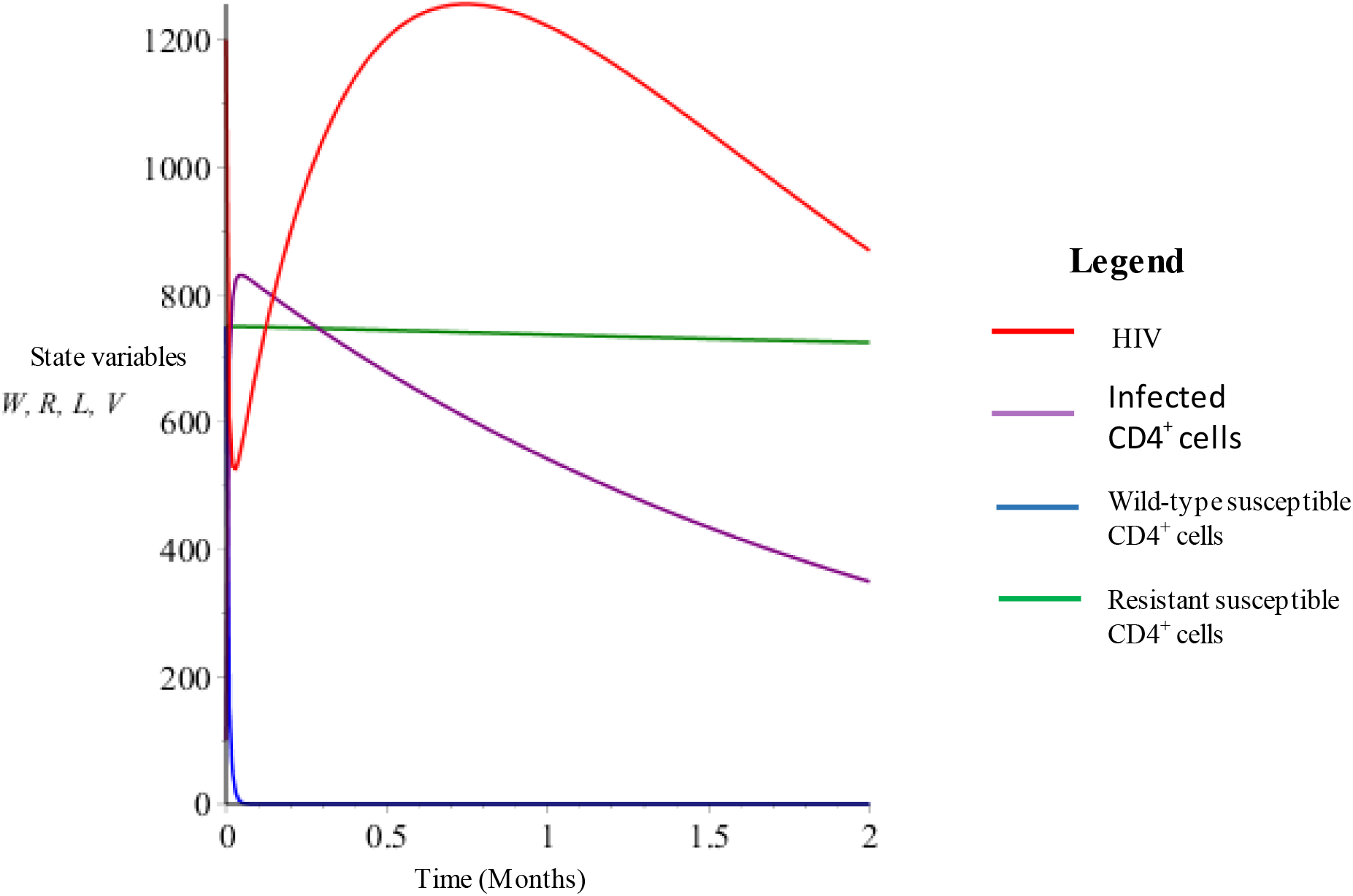
State variables vs time (β = 0.00001 resistant lymphocytes)

### 3.3 Changing a combination of state variables and force of infection of resistant lymphocytes

The simulated transmission dynamics up to this point have looked at the output of changing state variable or parameter initial values only. Transmission dynamics when initial value changes are made to parameters as well as state variables will be looked at next.

#### 3.3.1 Scenario 1: β = 0.005, V = 1200, W = 1200, R = 300, L = 100

The first scenario involves having an 80:20 population ratio of wild-type: resistant lymphocytes that are susceptible. This yields 300 resistant susceptibe lymphocytes and 1200 wild-type lymphocytes which is still a big enough pool of easily transmissible lymphocytes count to allow for a rapid peaking of the infected lymphocyte count (purple curve). The difference in transmission rate is maintained at resistant suscpeptible lymphocytes being 40 times less transmissible than the wild-type susceptible lymphocytes. The infected lymphocytes reach peak counts at about 4 months.

#### 3.3.2 Scenario 2: β = 0.005, V = 1200, W = 300, R = 1200, L = 100

Simply interchanging the relative population of wild-type versus resistant susceptible lymphocytes changes the dynamics dramatically until the peak of infected lymphocytes is attained at the 6-7 month time frame. The net result is an prolonged life expectancy of the lymphcytes and therefore of the host.

#### 3.3.3 Scenario 3: β = 0.005, V = 1200, W = 300, R = 1200, L = 100, Duration on the x-axis t = 10 months

The only difference between this Graph (Fig. 12) and the immediate previous one (Fig. 13) is the duration of simulation which has been chosen to be 10 months. In this scenario, it can be seen that as the resistant susceptible lymphocyte count is staying stable while the number of infected lymphocytes and that of virus drop precipitously.

**Figure 11:**
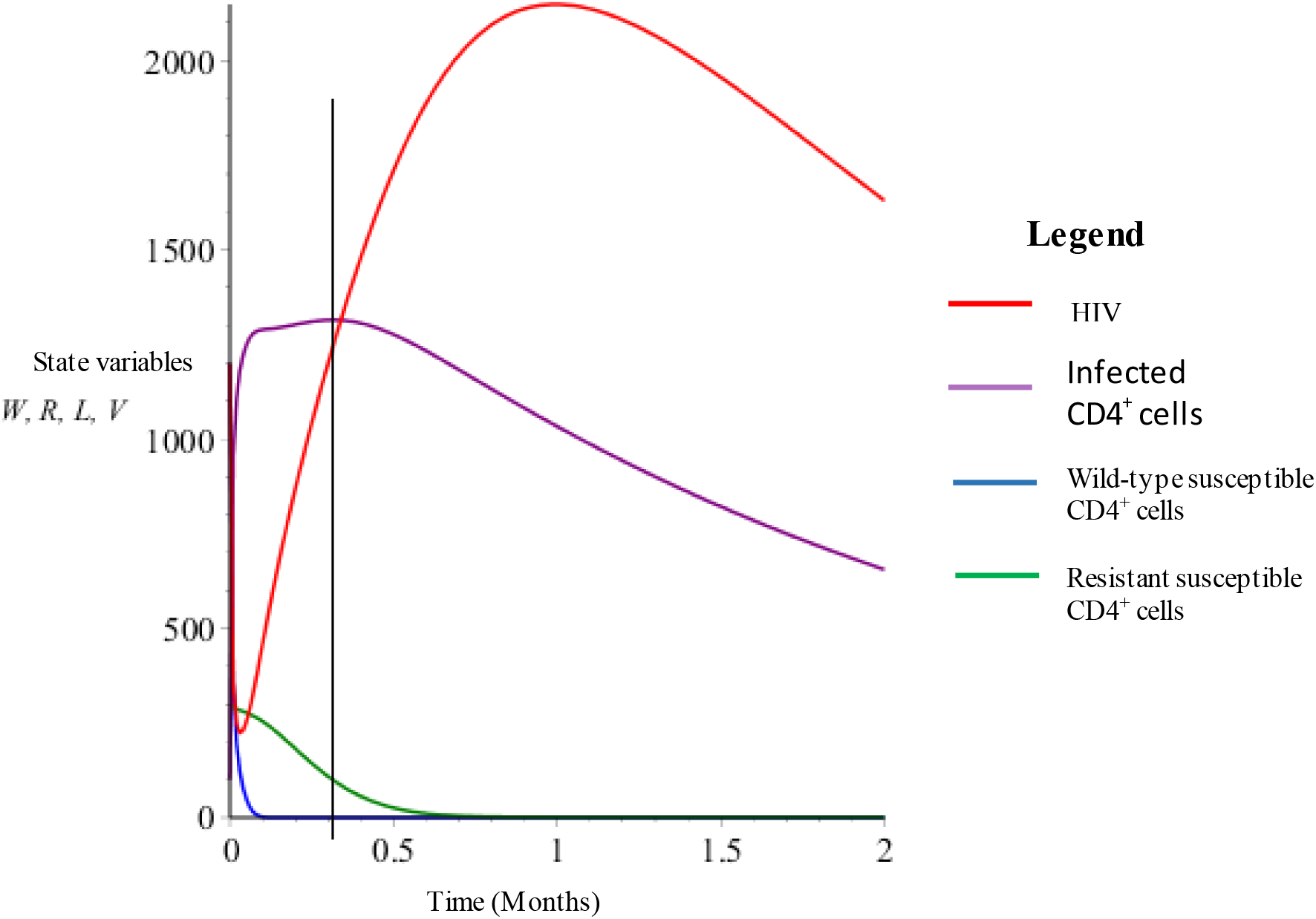
β = 0.005, V = 1200, W = 1200, R = 300, L = 100 (20% initially Resistant lymphocytes)

**Figure 12:**
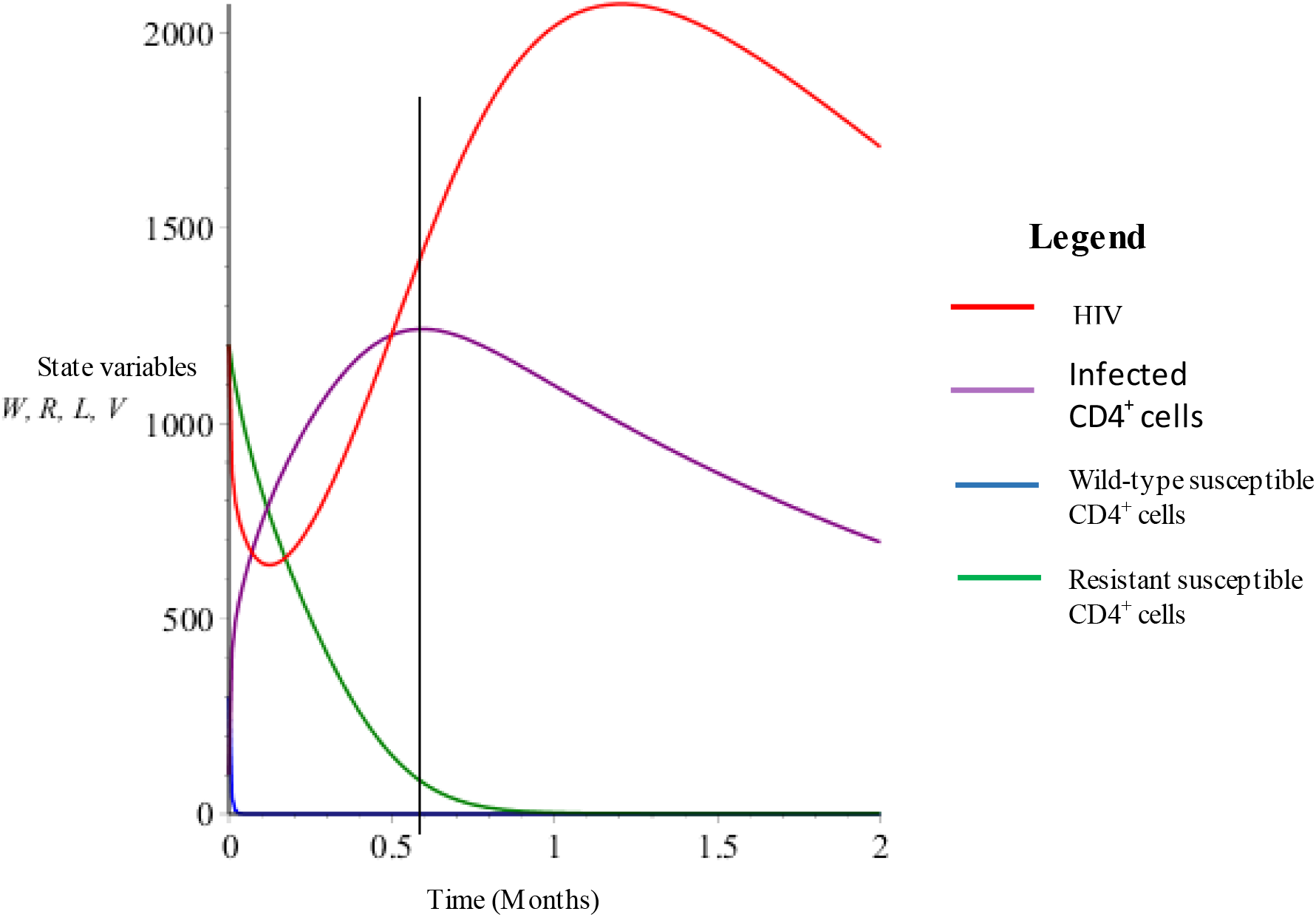
β = 0.005, V = 1200, W = 300, R = 1200, L = 100 (80% initially resistant lymphocytes)

**Figure 13:**
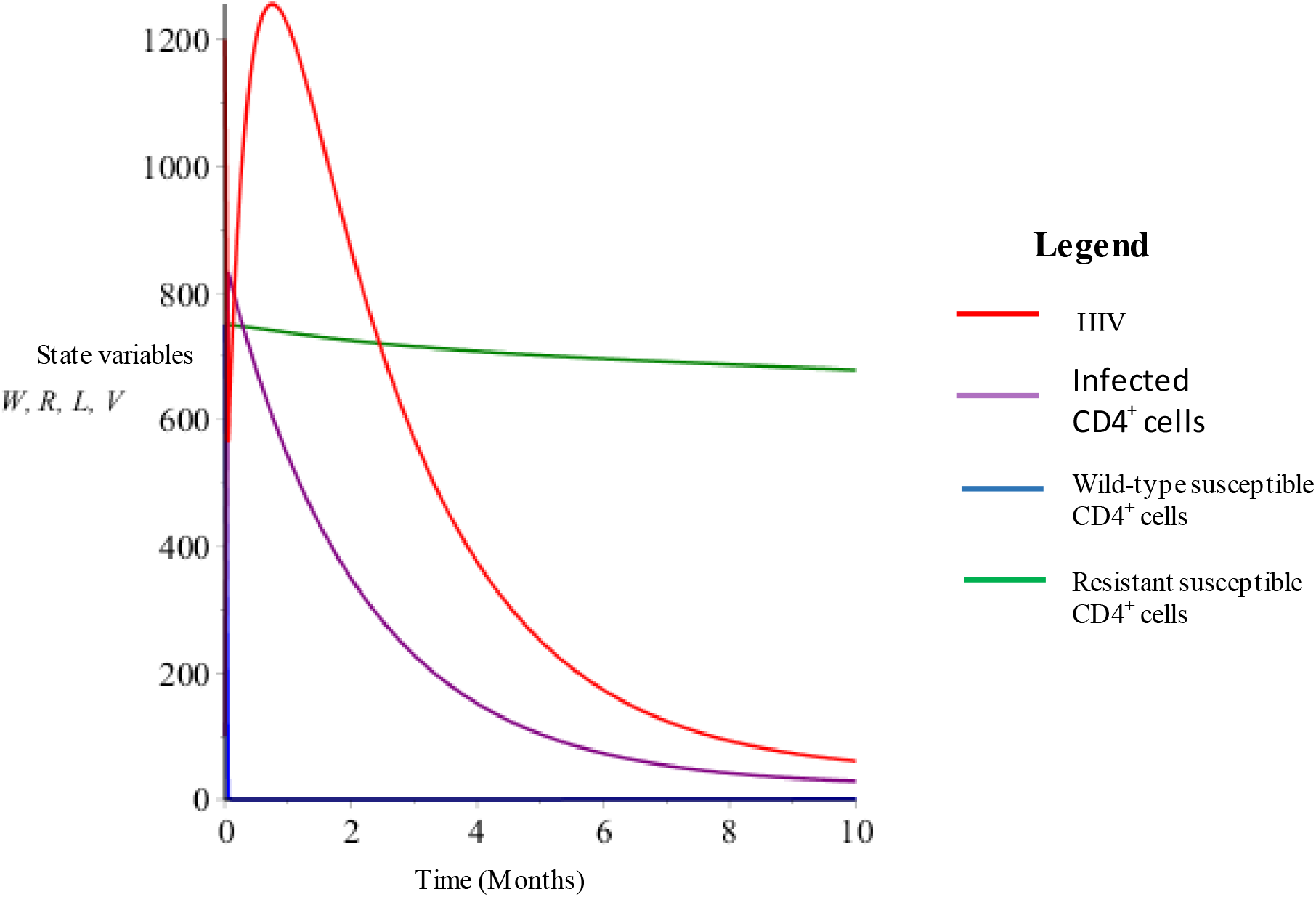
Same as Scenario 2 with a 10 month time line.

## 4 Discussions and Conclusions

Mathematical modeling findings from this project seem to support the, real world, HIV transmission dynamics that show varied resistant and susceptibility profiles of different hosts with some being relatively more resistant to HIV acquisition while others showing a more sensitive phenotype to HIV infection. Further work is necessary to display and characterize the probability distribution patterns of susceptible lymphocytes proportions in the in vivo ecosystem. As far as the specific dynamics of transmission of HIV are concerned, it is clear that great benefits can be gained by each host if the profile of relatively more resistant lymphocytes can be bolstered at the expense of the more wild-type variety. Many potential applications of this simulation could be produced from this mathematical model ranging from epidemiology, classification, investigation, diagnosis, monitoring, treatment, and even prognostication of both the HIV strains affecting individual hosts as well as the host’s response to the effects of infection.

Future directions of study may be aimed at identifying, describing, and profiling the different levels of resistance patterns of *CD4^+^* T-lymphocytes that play a critical role in the transmission dynamics of HIV.

